## Supplementary Information for "Galectin-9 Regulates The Threshold of B Cell Activation and Autoimmunity"

#### Supplementary Figure Legends

**Fig. S1 Nur77 reporter expression correlates with BCR internalization rate.** A) Representative histograms of Nur77 expression at 16 hours in WT (black) and Gal9KO (red) B cells stimulated with anti-IgM F(ab')$_2$ for 20 minutes, as indicated. B) Summary gMFI of Nur77 expression shown in A, mean and SEM. C) Correlation analysis of the internalization rate (k) of IgM-BCR over a 20 minute interval relative to the extent of BCR signal transduction as measured by the intensity of Nur77 expression, WT (left), Gal9KO (right). Data are representative of six biological replicates over two experiments. Statistical significance was assessed by Mann-Whitney ** $p \leq 0.01$

**Fig. S2 Gal9KO B cells have enhanced internalization of low affinity antigens.** A) Summary statistics of IgM internalization over 20 minutes as indicated, mean and SEM. B) Representative histograms of intracellular lysozyme levels at 20 minutes as indicated. Data are representative of nine biological replicates over three independent experiments.

**Fig. S3 Gal9KO B cells can escape anergy.** A) Representative plots of ASCs in the bone marrow of aged WT (left) and Gal9KO (right) mice. B) Proportion of ASC in the bone marrow of WT (black) and Gal9KO (red), as in A. C) Representative plots of Tfh cells in the spleen of WT (left) and Gal9KO (right) aged mice. D) Proportion of Tfh cells shown in C. E) Representative plots of FoxP3 expressing Tfr cells of Tfh cells in WT (left) and Gal9KO (right). F) Proportion of Tfr cells. G) Ratio of Tfh cells to regulatory Tfr cells in the spleen of aged mice. H) Representative plots of IgM high (hi) and IgM low (lo) populations in WT (left) and Gal9KO (right) MD4/ML5 mice. I) Proportion of $IgM^{hi}$ expressing cells shown in H. J) Representative histograms of p-Tyr levels in B cells 5 minutes post stimulation with HEL, as indicated. K) Summary gMFI of p-Tyr expression in J. L) Anti-HEL specific IgM titers in MD4 and Gal9KO-MD4 mice (left) and MD4/ML5 and Gal9KO-MD4/ML5 mice (right). Data show mean and SEM and are representative of nine to twelve biological replicates over at least three experiments. Statistical significance for B, D, F, G, I, and L was assessed by Mann-Whitney. Statistical significance for K was assessed by Kruskal-Wallis. ** $p \leq 0.01$, *** $p \leq 0.001$, **** $p < 0.0001$

**Fig. S4 B1-a cells from Gal9KO mice are activated at steady-state.** A) Representative histograms of CD69 expression on PerC cells in aged WT (black) and Gal9KO (red) mice, as indicated. B) Summary gMFI of CD69 expression shown in A. C) Representative histograms of MHCII expression on PerC cells in aged mice, as indicated. D) Summary gMFI of MHCII expression shown in C. E) Representative plots of B-1 cells in the thymus of aged WT (left) and Gal9KO (right) mice. F) Summary proportion of thymic B-1 cells of data shown in E. G) Total number of B-1 cells in the thymus in aged mice, as in F. Data represent ten biological replicates over three experiments. Statistical significance was assessed by Mann-Whitney *** $p \leq 0.001$, **** $p < 0.0001$

**Fig. S5 Gal9 does not alter BCR responses in B-1b cells.** A) Representative histograms of CD86 expression on B-1b cells from WT (black) and Gal9KO (red) mice stimulated with titrated concentrations of anti-IgM $F(ab')_2$. B) Summary proportion of CD86 expressing cells as in A. C) $EC_{50}$ of $F(ab')_2$ titration shown in A. D) Representative histograms of p-Tyr levels at 5 minutes in B-1b cells from WT (black, open) and Gal9KO (red, filled) mice, or WT B-1b cells treated with lactose (blue, open) and Gal9KO B-1b cells treated with rGal9 (orange, filled) stimulated with anti-IgM $(Fab')_2$ (left) and summary gMFI of p-Tyr level (right). E) Representative histograms of IgM detection on rGal9-coated beads incubated with lysates from B-1a (green), B-1b (white), and B-2 (purple) cells (left), and summary gMFI of CD5 enrichment on beads incubated with lysates (right). F) Reconstructed dSTORM images of IgM on the surface of B-1b cells. ROI (3 µm x 3 µm) used for analysis is expanded in zoom. Scale bars represent 1 µm. G) Quantification of IgM clustering tendency using Hopkins index. H) Quantification of IgM mean cluster area. Data in A-E are representative of nine biological replicates over three independent experiments. Data in G and I are representative of 30 ROIs acquired over at least three independent experiments. Data represent mean and SEM. Statistical significance for C, G and H was assessed by Mann-Whitney, statistical significance for D and E was assessed by Kruskal-Wallis. * $p \leq 0.05$, ** $p \leq 0.01$

**Fig. S6 Gal9 does not alter LPS responses in B-1b cells.** B1-b cells from WT (black) or Gal9KO (red) mice were stimulated with titrated concentrations of A) LPS, (B) CpG, (C) Imiquimod, or (D) Zymosan. Proportion of CD86 expressing B-1b cells shown on the left, and $EC_{50}$ on the right. E) Representative histogram of p-Tyr levels at 10 minutes in B-1b cells from WT (black, open) or Gal9KO (red, filled) mice, and WT B-1b cells treated with lactose (blue, open) or Gal9KO B-1b cells treated with rGal9 (orange, filled) and stimulated with LPS (left) and summary gMFI of p-Tyr levels (right). F) Reconstructed dual-dSTORM images of CD180 (magenta) and TLR4 (green) on the surface of B-1b cells. ROI (3 µm x 3 µm) used for analysis is expanded in zoom. Scale bars represent 1 µm. G) Quantification of TLR4 cluster area from dSTORM images. H) Quantification of the clustering tendency of TLR4 using the Hopkins index. I) Quantification of CD180 cluster area from dSTORM images. J) Quantification of CD180 clustering tendency using the Hopkins index. K) Frequency distribution of coordinate-based colocalization (CBC) between TLR4 and CD180. L) Mean CBC value per ROI. M) Mean distance between nearest neighbours. Data in A-E are representative of nine biological replicates over three independent experiments. Data in F-M are representative of 30 ROIs acquired over at least three independent experiments. Data represent mean and SEM. Statistical significance for A-D, G-I, J, L and M was assessed by Mann-Whitney, statistical significance for E was assessed by Kruskal-Wallis.

**Fig. S7 Gal9KO B-1a cells have enhanced activation to low LPS stimulation.** A) Representative histograms of CD86 expression on B-1a and B-1b cells from WT (black) or Gal9KO (red) mice injected i.p. with 1 µg/ml (top) or 0.05 µg/ml (bottom) of LPS (left), and proportion of CD86 expressing cells (right) as indicated. B) Fold increase in phosphorylcholine-specific IgM (left) and IgG (right) antibody titers in sera following LPS injection as in A. C) Representative histograms of IgG expression on splenic subcapsular sinus macrophages (left), summary fold increase in IgG gMFI following LPS injection (right). D) Representative histograms of IgM expression on splenic subcapsular sinus macrophages (left), summary fold increase in IgG gMFI following LPS injection (right). E) Representative histograms of apoptotic body detection on splenic subcapsular sinus macrophages (left), summary fold increase in apoptotic body gMFI following LPS injection and transfer of CFSE labeled apoptotic bodies (right). Data are representative of twelve biological replicates over three independent experiments. Data represent mean and SEM. Statistical significance was assessed by Mann-Whitney, * $p \leq 0.05$, ** $p \leq 0.01$

### Figure S1

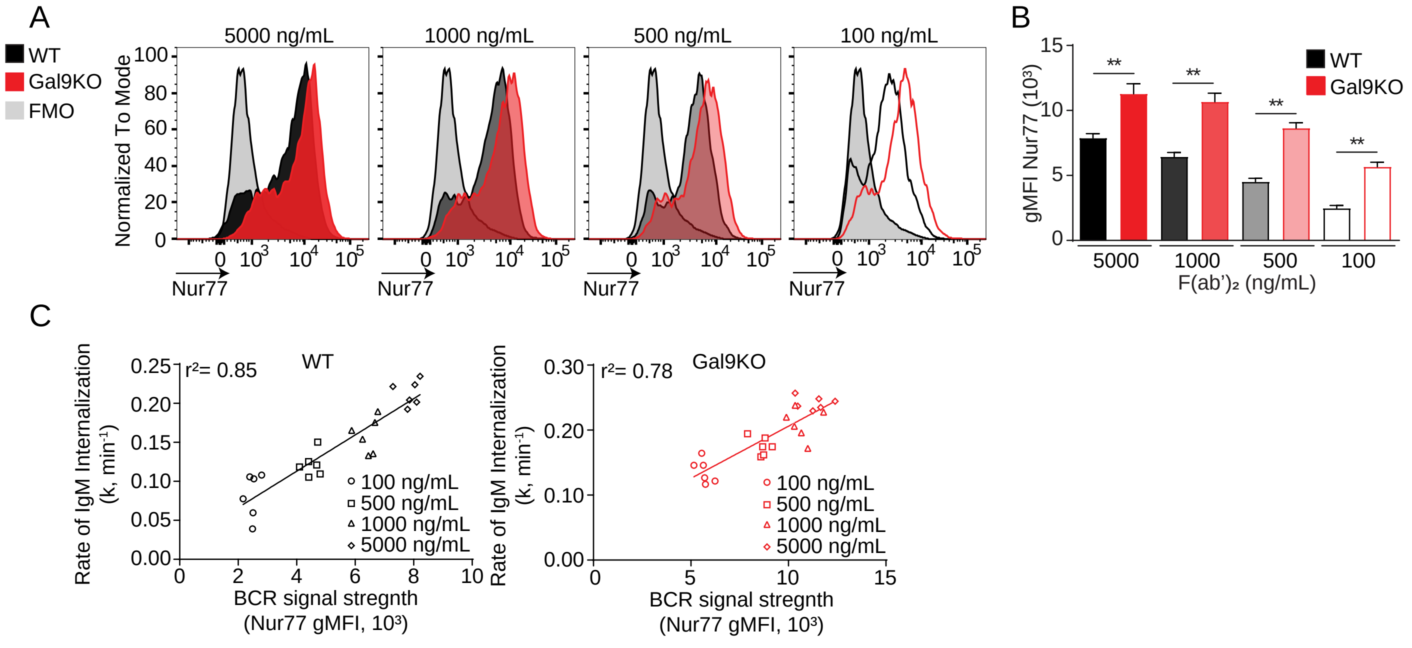

A
WT
Gal9KO
FMO
5000 ng/mL
1000 ng/mL
500 ng/mL
100 ng/mL
Normalized To Mode
Nur77
B
gMFI Nur77 (10³)
**
WT
Gal9KO
5000
1000
500
100
F(ab')₂ (ng/mL)
C
Rate of IgM Internalization (k, min⁻¹)
r²= 0.85
WT
100 ng/mL
500 ng/mL
1000 ng/mL
5000 ng/mL
BCR signal stregnth (Nur77 gMFI, 10³)
r²= 0.78
Gal9KO

### Figure S2

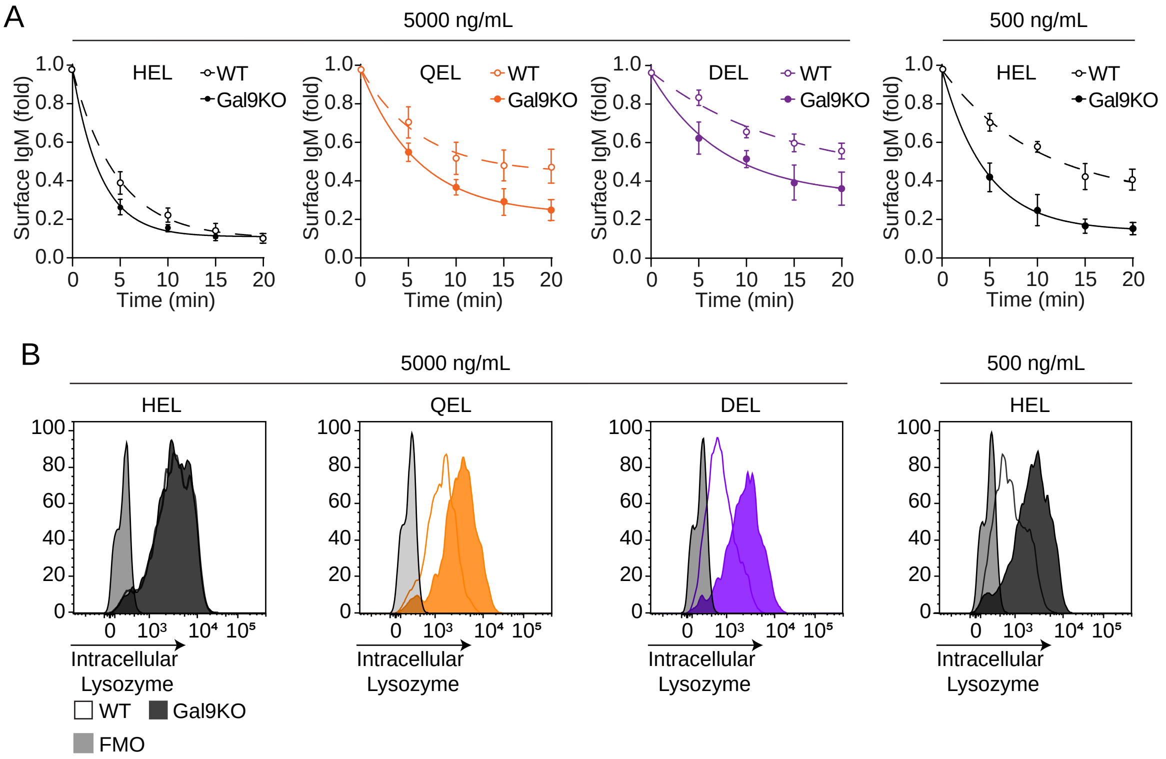

A
5000 ng/mL
500 ng/mL
HEL
QEL
DEL
HEL
WT
Gal9KO
Surface IgM (fold)
1.0
0.8
0.6
0.4
0.2
0.0
0
5
10
15
20
Time (min)
B
5000 ng/mL
500 ng/mL
HEL
QEL
DEL
HEL
100
80
60
40
20
0
0
10³
10⁴
10⁵
Intracellular Lysozyme
WT
Gal9KO
FMO

### Figure S3

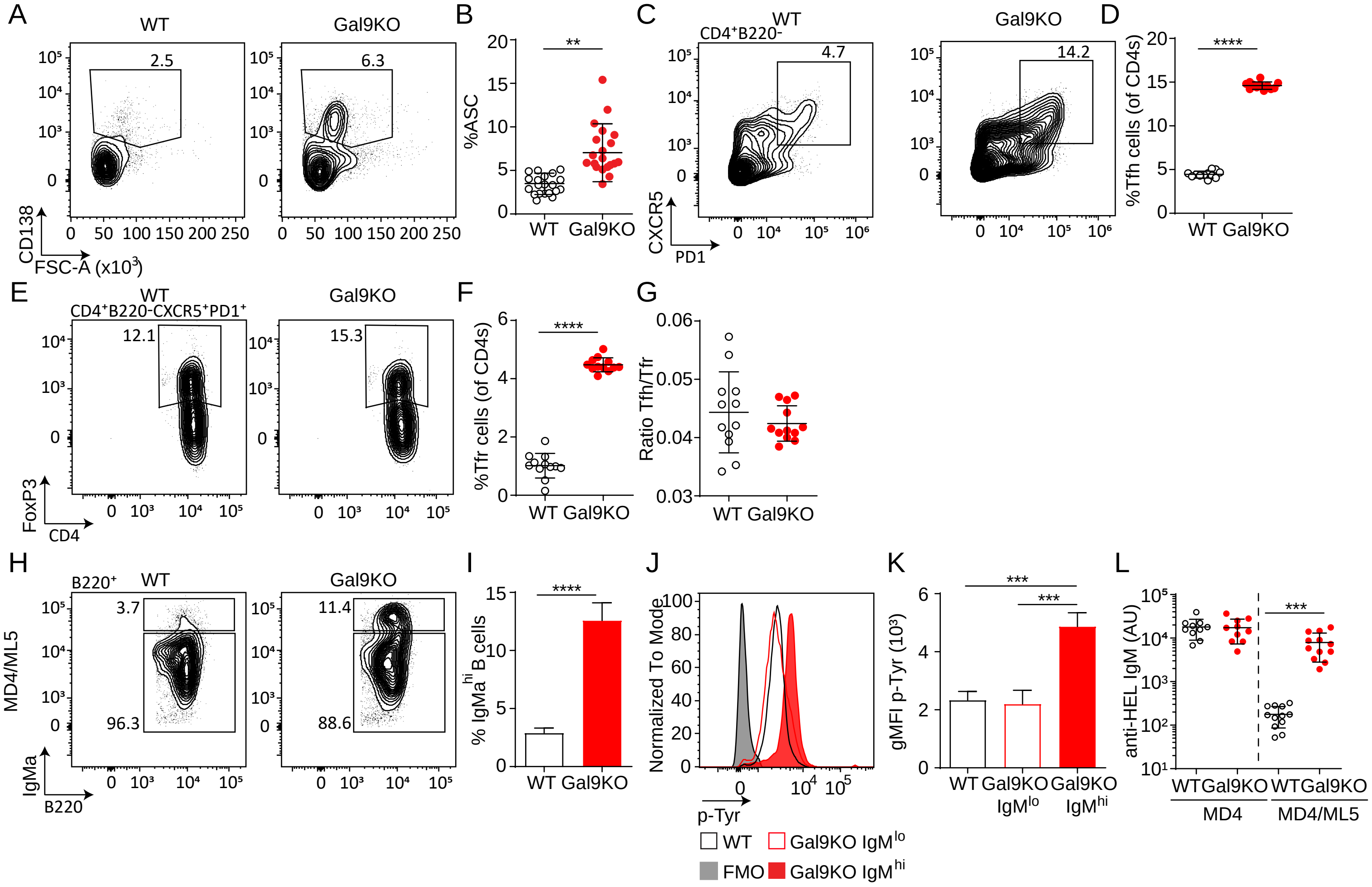

A
WT
Gal9KO
2.5
6.3
CD138
FSC-A (x10³)
B
%ASC
**
WT Gal9KO
C
CD4⁺B220-
4.7
14.2
CXCR5
PD1
D
%Tfh cells (of CD4s)
****
E
CD4⁺B220-CXCR5⁺PD1⁺
12.1
15.3
FoxP3
CD4
F
%Tfr cells (of CD4s)
G
Ratio Tfh/Tfr
H
B220⁺
3.7
96.3
11.4
88.6
MD4/ML5
IgMa
B220
I
% IgMa hi B cells
J
Normalized To Mode
p-Tyr
WT
Gal9KO IgMlo
FMO
Gal9KO IgMhi
K
gMFI p-Tyr (10³)
***
Gal9KO IgMlo
Gal9KO IgMhi
L
anti-HEL IgM (AU)
MD4
MD4/ML5

### Figure S4

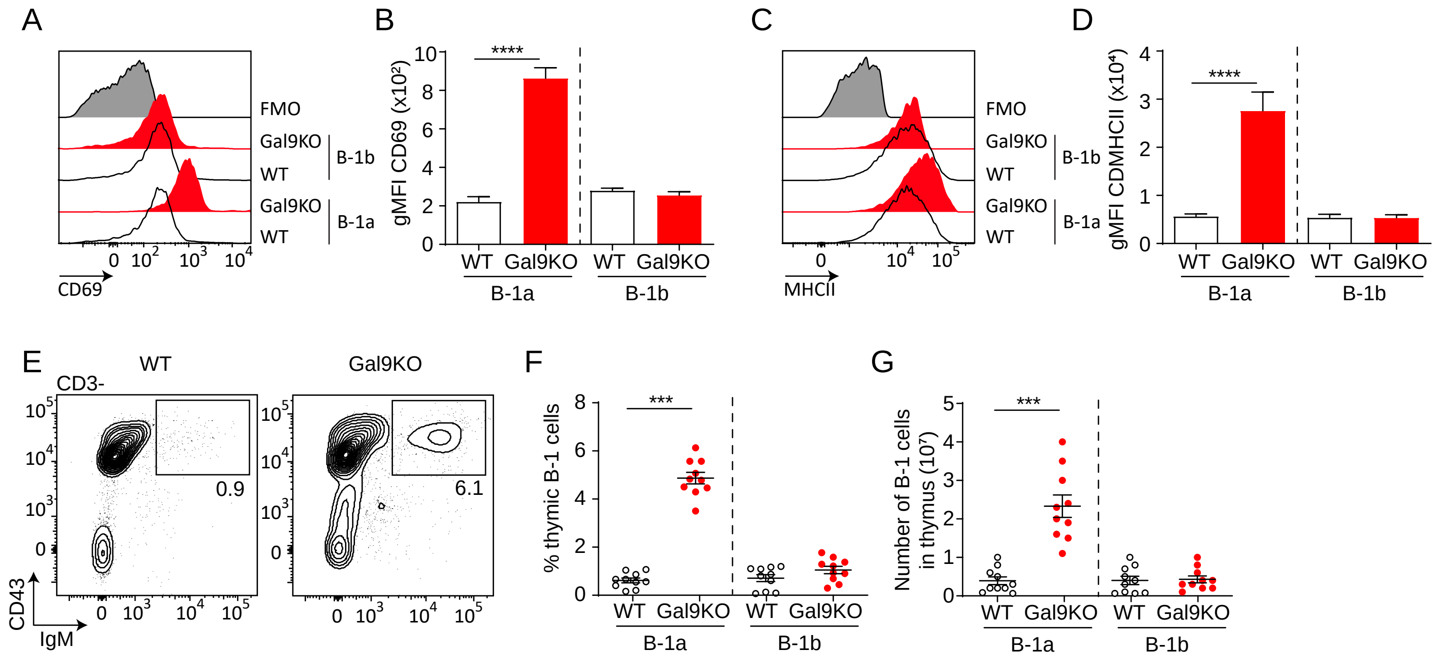
A
FMO
Gal9KO
WT
Gal9KO
WT
B-1b
B-1a
CD69
B
gMFI CD69 (x10²)
****
WT
Gal9KO
B-1a
B-1b
C
FMO
MHCII
D
gMFI CDMHCII (x10⁴)
E
WT
Gal9KO
CD3-
0.9
6.1
CD43
IgM
F
% thymic B-1 cells
***
G
Number of B-1 cells in thymus (10⁷)

### Figure S5

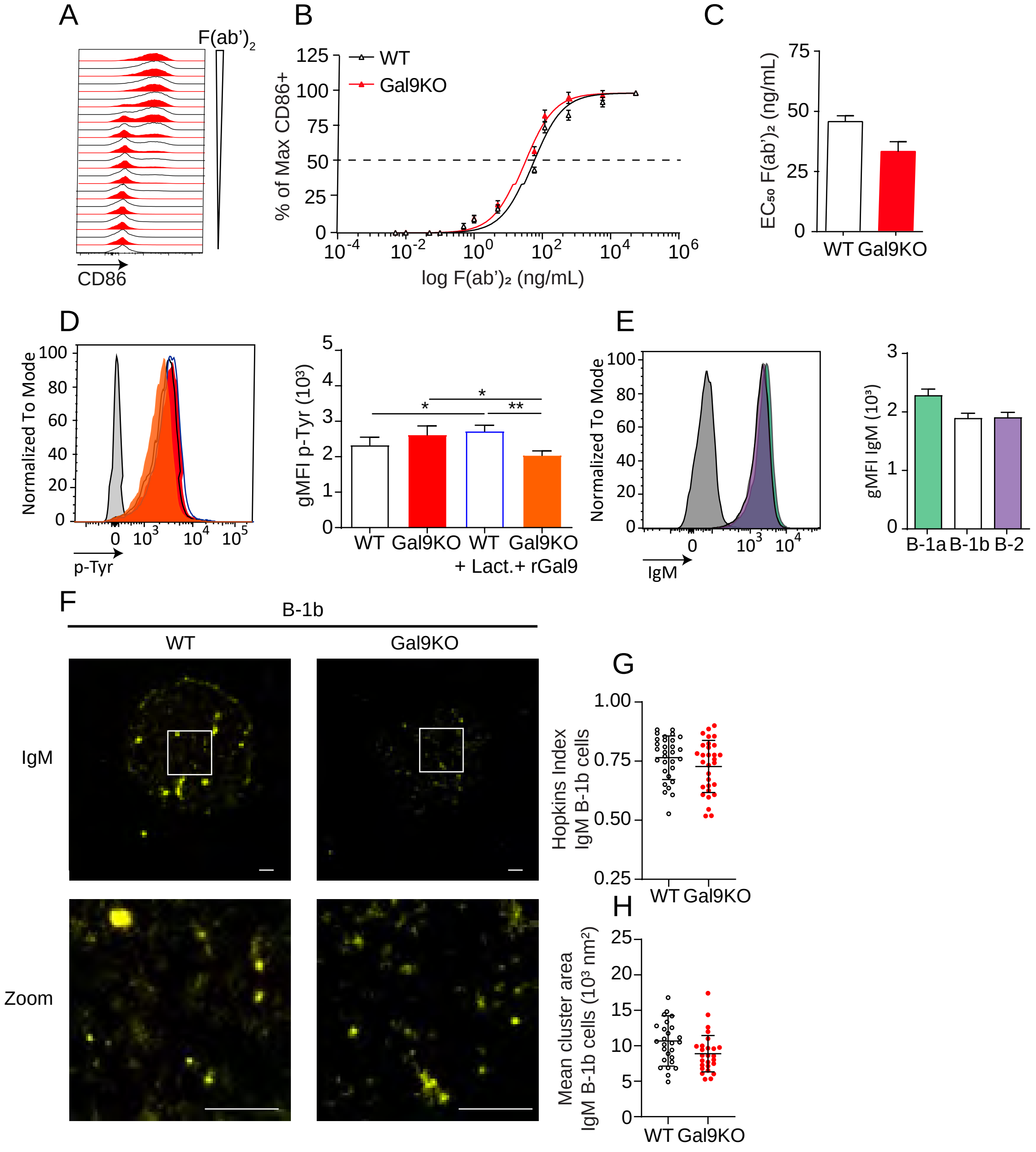
A
F(ab')2
CD86
B
WT
Gal9KO
% of Max CD86+
log F(ab')₂ (ng/mL)
C
EC₅₀ F(ab')₂ (ng/mL)
WT Gal9KO
D
Normalized To Mode
p-Tyr
gMFI p-Tyr (10³)
WT Gal9KO WT Gal9KO
+ Lact.+ rGal9
E
Normalized To Mode
IgM
gMFI IgM (10³)
B-1a B-1b B-2
F
B-1b
WT
Gal9KO
IgM
Zoom
G
Hopkins Index
IgM B-1b cells
WT Gal9KO
H
Mean cluster area
IgM B-1b cells (10³ nm²)
WT Gal9KO

### Figure S6

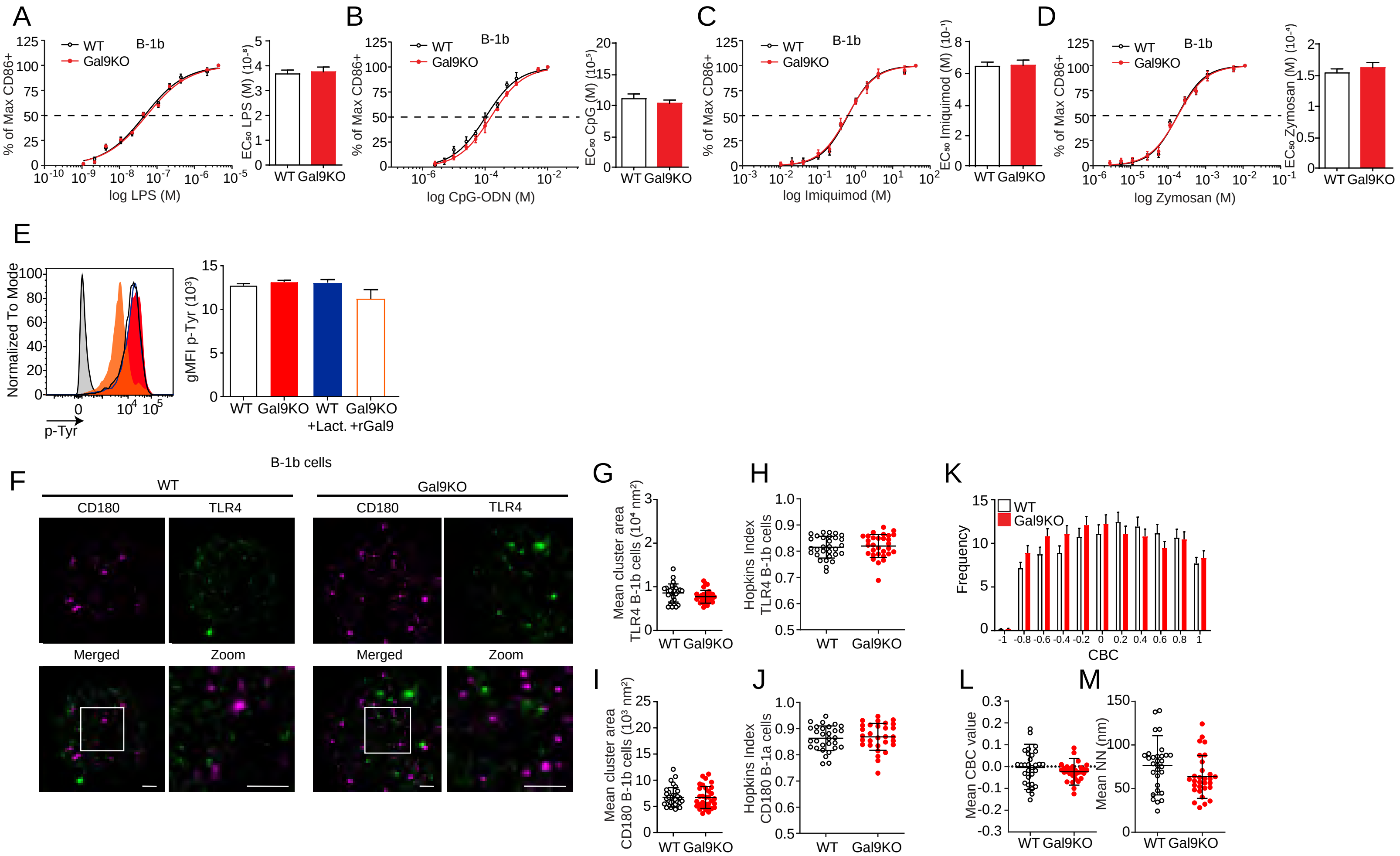
A
B
C
D
WT
Gal9KO
B-1b
% of Max CD86+
log LPS (M)
EC50 LPS (M) (10^-8)
log CpG-ODN (M)
EC50 CpG (M) (10^-5)
log Imiquimod (M)
EC50 Imiquimod (M) (10^-1)
log Zymosan (M)
EC50 Zymosan (M) (10^-4)
E
Normalized To Mode
p-Tyr
gMFI p-Tyr (10^3)
WT Gal9KO WT Gal9KO
+Lact. +rGal9
F
B-1b cells
WT
Gal9KO
CD180
TLR4
Merged
Zoom
G
Mean cluster area TLR4 B-1b cells (10^4 nm^2)
H
Hopkins Index TLR4 B-1b cells
K
Frequency
CBC
I
Mean cluster area CD180 B-1b cells (10^3 nm^2)
J
Hopkins Index CD180 B-1a cells
L
Mean CBC value
M
Mean NN (nm)

### Figure S7

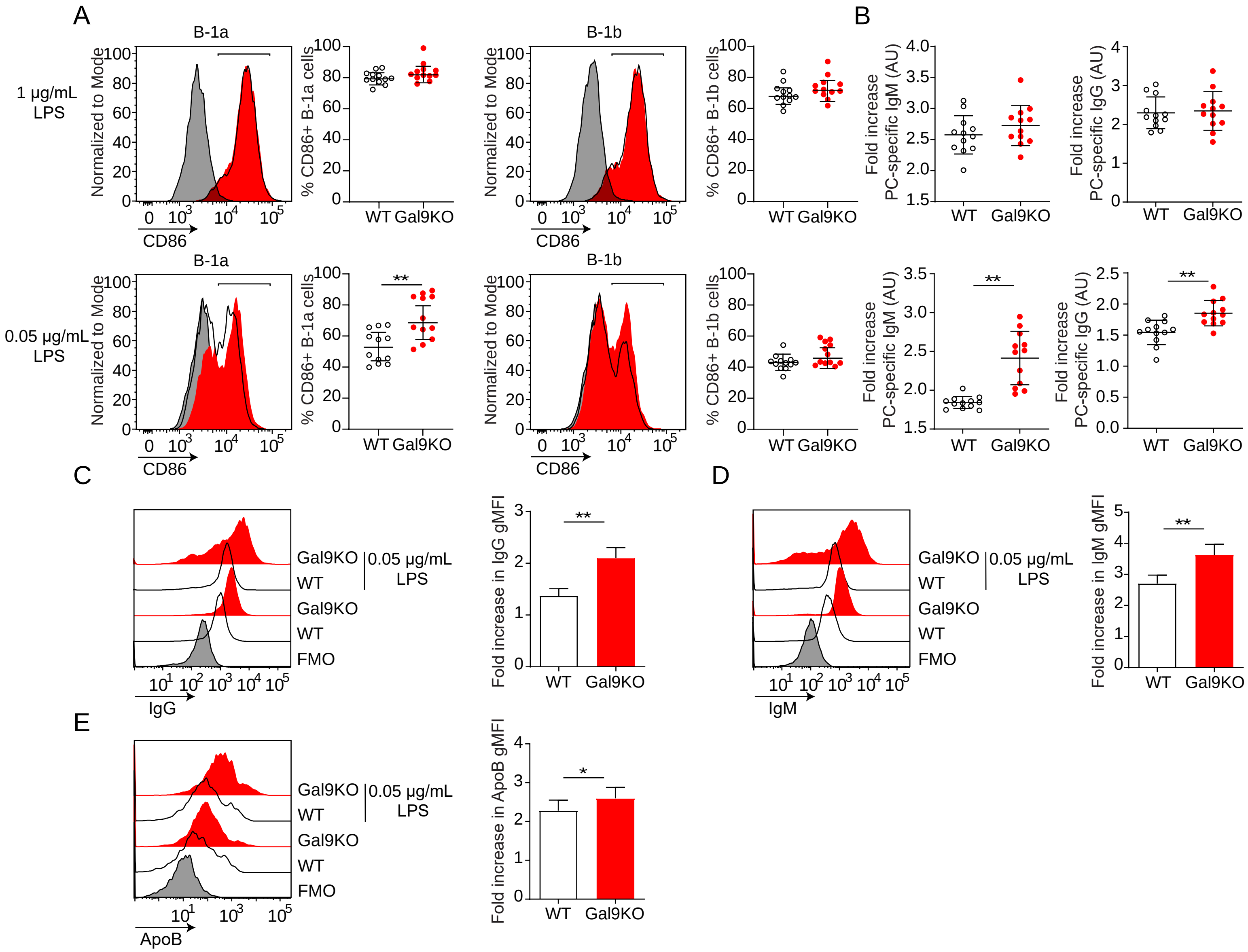

A
B
B-1a
B-1b
1 μg/mL LPS
0.05 μg/mL LPS
Normalized to Mode
CD86
% CD86+ B-1a cells
% CD86+ B-1b cells
WT
Gal9KO
Fold increase PC-specific IgM (AU)
Fold increase PC-specific IgG (AU)
**
C
D
E
Gal9KO
WT
FMO
0.05 μg/mL LPS
IgG
IgM
ApoB
Fold increase in IgG gMFI
Fold increase in IgM gMFI
Fold increase in ApoB gMFI
*
